# One-carbon metabolic enzymes are regulated during cell division and make distinct contributions to the metabolome and cell cycle progression in *Saccharomyces cerevisiae*

**DOI:** 10.1101/2022.10.25.513769

**Authors:** Staci E. Hammer, Michael Polymenis

**Affiliations:** Department of Biochemistry and Biophysics, Texas A&M University, College Station, TX 77843

**Author notes:** **CORRESPONDENCE:** Michael Polymenis, 300 Olsen, Blvd, 2128 TAMU, College Station, TX 77843.

**Keywords:** *ADE17*, *SHM2*, *CHO2*, cell size, nuclear division, phosphatidylcholine

## Abstract

Enzymes of one-carbon metabolism play pivotal roles in proliferating cells. They are involved in the metabolism of amino acids, nucleotides, and lipids and the supply of all cellular methylations. However, there is limited information about how these enzymes are regulated during cell division and how cell cycle kinetics are affected in several loss-of-function mutants of one-carbon metabolism. Here, we report that the levels of the *S. cerevisiae* enzymes Ade17p and Cho2p, involved in the *de novo* synthesis of purines and phosphatidylcholine, respectively, are cell cycle-regulated. Cells lacking Ade17p, Cho2p, or Shm2p (an enzyme that supplies one-carbon units from serine) have distinct alterations in size homeostasis and cell cycle kinetics. Loss of Ade17p leads to a specific delay at START, when cells commit to a new round of cell division, while loss of Shm2p has broader effects, reducing growth rate. Furthermore, the inability to synthesize phosphatidylcholine *de novo* in *cho2Δ* cells delays START and reduces the coherence of nuclear elongation late in the cell cycle. Loss of Cho2p also leads to profound metabolite changes. Besides the expected changes in the lipidome, *cho2Δ* cells have reduced levels of amino acids, resembling cells shifted to poorer media. These results reveal the different ways that one-carbon metabolism allocates resources to affect cell proliferation at multiple cell cycle transitions.

## INTRODUCTION

One-carbon (1C) metabolism encompasses the chemical reactions that move and use single-carbon functional groups. At the heart of 1C metabolism is tetrahydrofolate (THF; see Figure 1). THF carries activated 1C units at various oxidation states (Appling *et al*.2019). The most reduced form is methyl-THF, which donates the methyl group only to homocysteine, forming the C-S bond of methionine, and then, through S-adenosylmethionine (AdoMet, SAM), supplying all cellular methylations, including on phosphatidylcholine (PC) and histones (Figure 1). The 1C unit of methylene-THF (CH_2_-THF) forms new C-C bonds in the synthesis of dTMP and the interconversions involving serine and glycine. Formyl-THF (CHO-THF) carries the most oxidized 1C units, forming C-N bonds in the synthesis of purines.

**FIGURE 1.**
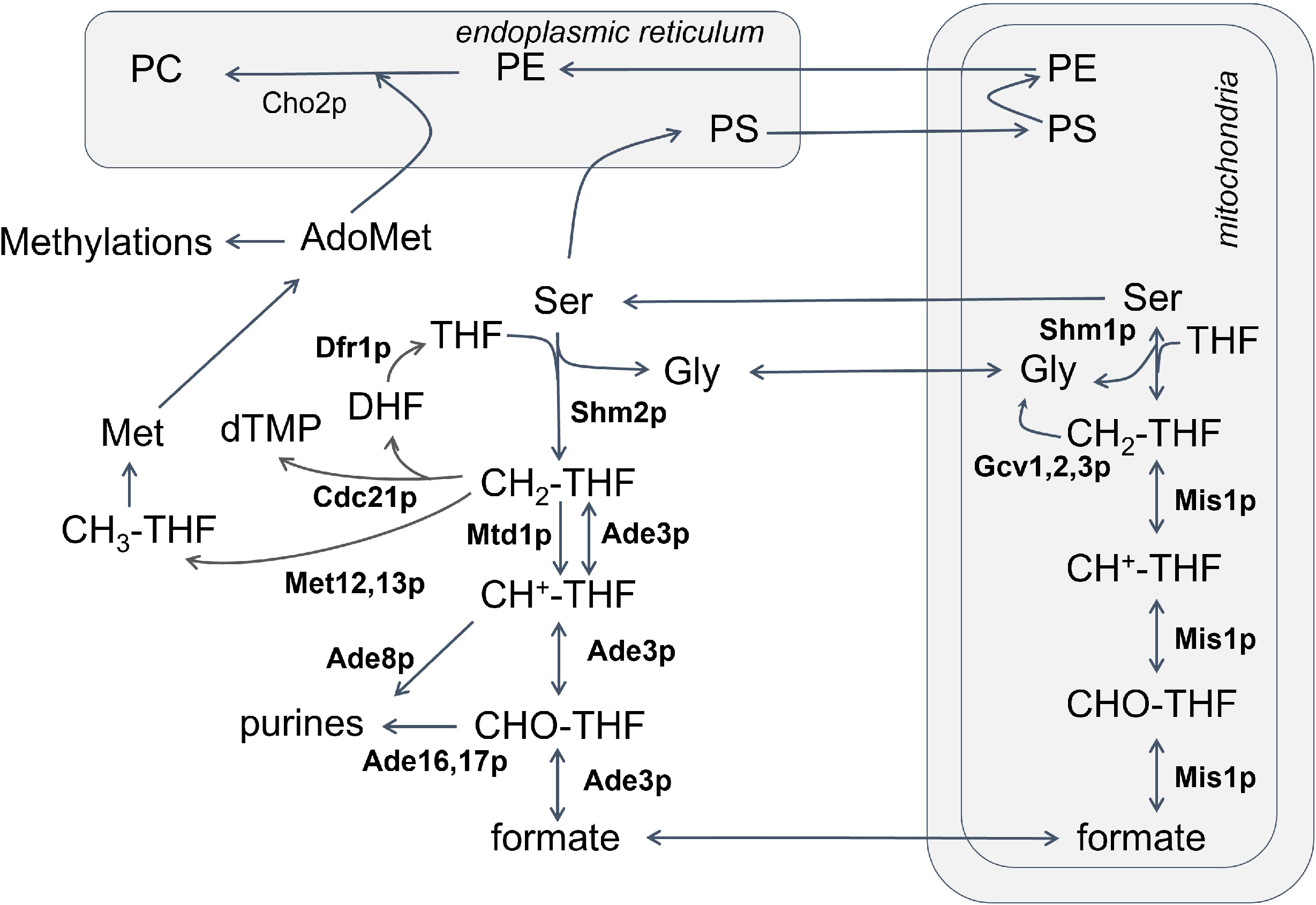
Schematic of 1C reactions. Enzymes involved in folate interconversions in *S. cerevisiae* are shown in bold. The pathway for the *de novo* synthesis of phosphatidylcholine, the major consumer of 1C units in the cell, and the step catalyzed by Cho2p are also indicated.

1C pathways are directly involved in the metabolism of amino acids and the synthesis of purines, thymidylate, and phospholipids (West *et al*. 1996; Fox and Stover 2008; Ducker and Rabinowitz 2017). As a result, 1C outputs govern vital cellular processes, including genome replication and maintenance (through nucleotide synthesis), response to oxidative stress (through glutathione synthesis), gene expression (through methylation of DNA and histones), among others (Fox and Stover 2008; Ducker and Rabinowitz 2017). 1C pathways receive enormous attention in diseases, especially cancer (Locasale 2013; Labuschagne *et al*. 2014; Rosenzweig *et al*. 2018; Reina-Campos *et al*. 2019). For example, during dTMP synthesis by thymidylate synthase (Cdc21p in *S. cerevisiae*), the methylene-THF coenzyme is the 1C unit source and the reducing power, getting oxidized in the process to DHF. To maintain the THF levels needed in the cells, dihydrofolate reductase (Dfr1p in *S. cerevisiae*) reduces DHF back to THF (Figure 1). Methotrexate is one of the oldest chemotherapeutics and a potent and specific inhibitor of dihydrofolate reductase (Williams *et al*. 1979), leading to THF deficiency, accounting for its antiproliferative properties. In a recent extensive transcriptomic profiling of 1,454 metabolic enzymes across 1,981 tumors among 19 cancer types, 1C enzymes had the highest score for being consistently over-expressed compared to non-cancerous tissue samples (Nilsson *et al*. 2014).

Yet, despite the significance of 1C metabolism for cell proliferation and the vast literature dealing with 1C-based anti-proliferative interventions, how the activity of 1C enzymes is regulated in the cell cycle of normal cells is not well understood. Changes in the levels of 1C enzymes are expected to change 1C metabolic outputs because the levels of 1C enzymes exceed by several-fold the folate metabolites, and the enzymes compete with each other for the available folate pools (Fox and Stover 2008; Lan *et al*.2018). In mammalian cells, some reports suggested that mRNAs encoding 1C enzymes may be periodic in the cell cycle (summarized in (Lan *et al*. 2018)). However, based on aggregate transcriptomic datasets, only the levels of *DHFRL1* (encoding mitochondrial dihydrofolate reductase) and *TYMS* (encoding thymidylate synthetase) change significantly in the cell cycle, peaking in the S phase (Santos *et al*. 2015). Likewise, in the budding yeast *Saccharomyces cerevisiae*, only the mRNAs encoding thymidylate synthase (*CDC21*) and a subunit of the mitochondrial glycine decarboxylase complex (*GCV2*) are periodic in the cell cycle, peaking at the G1/S transition, and the G2 phase, respectively (Santos *et al*. 2015). In addition to transcription, other possible layers of temporal control of 1C enzymes in the cell cycle include translational, post-translational, and differential subcellular localization. Indeed, in mammalian cells, serine hydroxymethyltransferase isoforms (SHMT2α and SHMT1) translocate in the nucleus during the S phase, presumably to provide 1C units needed for thymidylate synthesis and DNA replication (Woeller *et al*. 2007; Anderson and Stover 2009; Anderson *et al*.2012; Lan *et al*. 2018). In HeLa cells blocked with hydroxyurea in the S phase, SHMT1 protein levels were elevated without concomitant changes in *SHMT1* mRNA levels (Anderson *et al*. 2012).

In budding yeast, translational control could impose temporal control of 1C metabolism in the cell cycle. The translational efficiency of mRNAs encoding several 1C enzymes was altered in the cell cycle based on ribosome profiling experiments previously reported in ribosomal protein mutants (Maitra *et al*. 2020). These mRNAs included *ADE17* (ATIC in humans; encoding both a 5-aminoimidazole-4-carboxamide ribonucleotide transformylase and inosine monophosphate cyclohydrolase activities), *SHM2* (SHMT1 in humans; encoding the cytoplasmic serine hydroxymethyltransferase), and *CHO2* (PEMT in humans; encoding a phosphatidylethanolamine (PE) methyltransferase).

The above enzymes play critical roles in 1C metabolism. Ade17p controls the use of 1C units, in the form of CHO-THF, for purine synthesis (Tibbetts and Appling 2000). The cytoplasmic serine hydroxymethyltransferase (Shm2p in yeast) is a key metabolic switch in 1C metabolism (Piper *et al*. 2000; Herbig *et al*. 2002). Shm2p controls the input of carbon in the pathway from serine. Shm2p also contributes to the cytoplasmic CH2-THF pool towards dTMP and methylations, or away from them, to make more serine for phospholipids (from glycine and 1C units) (Piper *et al*. 2000; Herbig *et al*. 2002). It is serine, but not glycine, that supports 1C metabolism in cancer cells (Labuschagne *et al*. 2014). Serine supplementation, through 1C metabolism, also regulates chronological lifespan in budding yeast (Enriquez-Hesles *et al*. 2021). Although Cho2p is not a folate-dependent enzyme, it catalyzes the first step in the conversion of PE to PC during the methylation pathway of *de novo* PC biosynthesis. It has been reported that PC synthesis is the major consumer of 1C-derived methyl groups in yeast (Ye *et al*. 2017) and likely in humans (Stead *et al*. 2006). Cho2p has been proposed to control the flow toward phospholipid synthesis but away from histone methylation (Ye *et al*. 2017). However, it is not known if the levels of any of the above enzymes change in the cell cycle of unperturbed cells in any system. Furthermore, despite the numerous ways 1C metabolism could impact cell division, besides DNA replication, if and how specific 1C enzymes might control other cell cycle transitions is unknown.

Here, we report that the levels of Ade17p and Cho2p, and to a lesser extent that of Shm2p, are periodic in the budding yeast cell cycle. We also present an analysis of size homeostasis, cell cycle kinetics, and metabolic outputs in cells lacking any of these enzymes. These parameters were altered in the mutants we examined but in distinct ways in each mutant. Lastly, our data argue that Cho2p and *de novo* synthesis of PC are required to establish the proper kinetics of nuclear division. Overall, this work provides new and important information on the abundance and roles of 1C enzymes during cell division.

## MATERIALS AND METHODS

A Reagent Table is in the Supplementary Files. Where known, the Research Resource Identifiers (RRIDs) are shown in the Reagent Table.

### Strains and media

All the strains used in this study are shown in the Reagent Table. For most experiments, the cells were cultivated in the standard, rich, undefined medium YPD (1%^w^/_v_ yeast extract, 2%^w^/_v_ peptone, 2%^w^/_v_ dextrose), at 30° (Kaiser *et al*. 1994). To generate the *cho2Δ* haploid strain, we sporulated and dissected the commercially available homozygous diploid *cho2Δ/cho2Δ* strain (see Reagent Table). For the experiments with *cho2Δ* cells, we also used synthetic minimal media (SMM), containing 0.17%^w^/_v_ yeast nitrogen base without amino acids, 0.5%^w^/_v_ ammonium sulfate, 2%^w^/_v_ glucose, the required amino acid auxotrophies at standard concentrations (Kaiser *et al*. 1994) and, if indicated, 1 mM choline chloride.

Single gene homozygous deletion strains, lacking *ADE17, SHM2*, or *CHO2*, were commercially available (see Reagent Table). Their genotype was validated by PCR, to confirm that the gene of interest was absent and replaced by the appropriate marker. To construct the *ADE17-TAP* strain (see Reagent Table), a PCR product was generated using plasmid pBS1539 as a template (Puig *et al*. 2001), with primers Ade17-TAP-FWD and Ade17-TAP-REV, and used to transform strain BY4741. *SHM2-TAP* and *CHO2-TAP* strains were commercially available (see Reagent Table).

### Multiple correspondence analysis (MCA)

Data collection and analyses were done as described previously (Bermudez *et al*.2020; Polymenis 2020). All the relevant data are in File S2.

### Immunoblot analysis

For protein surveillance, protein extracts were made as described previously (Amberg *et al*. 2006), and resolved on 12% Tris-Glycine SDS-PAGE gels, unless indicated otherwise. To detect TAP-tagged proteins with the PAP reagent (used at 1:4,000 dilution), we used immunoblots from extracts of the indicated strains, as we described previously (Blank *et al*. 2017, 2020; Maitra *et al*. 2020, 2022). Loading was measured with an anti-Pgk1p primary antibody (at 1:2,000; abcam, Cat#:ab38007), followed by a secondary antibody (at 1:1,500; Jackson Immunoresearch Laboratories, Alexa Fluor^®^ 488 AffiniPure Goat Anti-Mouse IgG (H+L)). Imaging and quantification was done as described previously (Blank *et al*. 2017, 2020; Maitra *et al*. 2020, 2022).

### Centrifugal elutriation, cell size and DNA content measurements

All methods have been described previously (Hoose *et al*. 2012; Soma *et al*. 2014). Briefly, after early G1 cells were collected, they were monitored at regular time intervals for cell size, budding, or DNA content. Samples were also assayed in downstream procedures, such as nuclear staining, as described in the relevant sections.

### Metabolite profiling

The untargeted, primary metabolite, biogenic amine, and complex lipid analyses were done at the NIH-funded West Coast Metabolomics Center at the University of California at Davis, according to their established mass spectrometry protocols, as we described previously (Blank *et al*. 2020; Maitra *et al*. 2020, 2022). The analysis was done from six independent samples in each case. The raw data for the primary metabolite measurements are in File S3. The raw data for the primary amine measurements are in File S4. The raw data for the complex lipid measurements are in File S5. To identify significant differences in the comparisons among the different strains and media, based on peak intensities from each metabolite, we used the robust bootstrap ANOVA, as described below. Detected species that could not be assigned to any compound were excluded from the analysis. In the complex lipid analyses we also excluded species that are not present in yeast, such as those containing polyunsaturated fatty acid side-chains.

### Fluorescence microscopy

To visualize the nuclear envelope, we followed the same procedures we described previously (Maitra *et al*. 2022). Briefly, when the nucleus is near-spherical, the ratio of the long to short nuclear axes is 1. However, as the nucleus starts to expand, the ratio increases, signifying the elongation of the nuclear envelope (Figure 6).

### Histone methylation

Overnight cultures in YPD or SMM media of the indicated strains were diluted 1:100. To supplement SMM with choline, choline chloride was added at 1 mM, 2 h after dilution. After several hours, cells were harvested at 1E+07 cells/mL. Protein extracts were prepared and analyzed by SDS-PAGE and immunoblotting as described above. Primary antibodies were used to detect histone H3 (at 1:1,000 dilution; Abcam, Cat#ab1719) and H3K4me3 (at 1:2,000 dilution; Epicypher, Cat#13-0041), followed by an anti-rabbit secondary antibody (used at 1:2,000 dilution). Pgk1p was detected as described above. Protein bands were quantified with ImageJ. H3 and H3K4me3 signals were normalized to the respective Pgk1p bands. Methylation levels were then calculated by the ratio of the normalized H3 levels over the normalized H3K4me3 levels. The quantification is in Figure S3 and all the immunoblot source data are in Figure S4.

### Statistical analysis, sample-size and replicates

For sample-size estimation, no explicit power analysis was used. All the replicates in every experiment shown were biological ones, from independent cultures. A minimum of three biological replicates was analyzed in each case, as indicated in each corresponding figure’s legends. The robust bootstrap ANOVA was used to compare different populations via the t1waybt function, and the posthoc tests via the mcppb20 function, of the WRS2 R language package (Wilcox 2011; Mair and Wilcox 2016). We also used non-parametric statistical methods, as indicated in each case. The Kruskal-Wallis and posthoc Nemenyi tests were done with the posthoc.kruskal.nemenyi.test function of the PMCMR R language package. No data or outliers were excluded from any analysis.

## RESULTS

### The levels of Ade17p and Cho2p peak late in the cell cycle

The steady-state levels of *ADE17, SHM2*, or *CHO2* transcripts do not change in the cell cycle (Spellman *et al*. 1998; Santos *et al*. 2015; Blank *et al*. 2017, 2020). However, ribosome profiling experiments suggested that the translational efficiency of those transcripts changes in dividing cells of ribosomal protein mutants (Maitra *et al*. 2020). To monitor the levels of the corresponding proteins in the cell cycle, we used otherwise wild type, haploid strains carrying alleles encoding C-terminal, TAP-tagged versions of these enzymes (Puig *et al*. 2001), expressed from their endogenous locations in the genome. To maintain the physiological coupling between cell growth and division, we used centrifugal elutriation to prepare growing, synchronous cultures (Aramayo and Polymenis 2017). In *S.cerevisiae*, budding marks the initiation of cell division, and in daughter cells, the cell size is a proxy for cell cycle position (Hartwell and Unger 1977; Johnston *et al*. 1977). Newborn daughter cells in early G1 were sampled at regular intervals as they progressed in the cell cycle, recording their size and budding (Figure 1).

The levels of Ade17p-TAP increased markedly (~10-fold) from the beginning to the end of the cell cycle (Figure 1, left panels). Likewise, Cho2p-TAP levels were higher (~4-fold) late in the cell cycle, when the cells were large and budded (Figure 1, right panels). In contrast, Shm2p-TAP levels were the highest in the early G1 phase and then declined slightly (<2-fold) as the cells progressed in the cell cycle (Figure 1, middle panels). These results show that the levels of Ade17p and Cho2p are periodic in the cell cycle. They are also consistent with the notion that translational control of these gene products contributes to their oscillation in dividing cells.

### Cell cycle phenotypes of folate-dependent enzymes

Besides *cdc21* (encoding thymidylate synthase), identified in the classic *c*ell *d*ivision *c*ycle screen and shown to arrest in the S phase (Hartwell *et al*. 1974), there is limited information about cell cycle-related phenotypes of other folate-dependent enzymes. We generated a complete matrix of all the loss-of-function phenotypes associated with the 15 genes encoding enzymes that catalyze folate-dependent reactions (shown in bold in Figure 1; File S1) from the available data on the Saccharomyces Genome Database (Cherry *et al*. 2012). Sixty-six loss-of-function phenotypes are associated with the 15 folate-dependent enzymes (File S2/Sheet1). The most common phenotypes, reported for at least 6 of the 15 genes, are rather generic, reflecting altered competitive fitness in some conditions and chemical resistance. In contrast, only three genes have been associated with altered cell cycle progression: *CDC21* (arrest in S phase), *DFR1*(abnormal S phase), and *MIS1* (G1 delay). To identify any patterns that may underlie the observed phenotypes of 1C mutants, we applied multiple correspondence analysis (analogous to principal component analysis, but for categorical data) to define and group those phenotypes and the genes that may be primarily responsible for them (Bermudez *et al*. 2020; Polymenis 2020).

The 66 different phenotypes can be reduced to 14 dimensions, with just three dimensions accounting for 68% of the observed variance (Figure 3A; File 2/Sheet2). The major phenotypes contributing to Dimension 1 were abnormal cell morphology, meiotic arrest, lack of sporulation, cell cycle arrest, increased chitin deposition, decreased resistance to toxins, and petite colony formation (File S2/Sheet3). By far, the primary gene driving this grouping was *CDC21*(75% contribution; see Figure 3B and File S2/Sheet4). The major phenotypes contributing to Dimension 2 appear unrelated to cell cycle progression (e.g., abnormal endoplasmic reticulum morphology, increased RNA accumulation, and stress resistance; see File S2/Sheet3). The classification was driven chiefly by *gcv3* phenotypes (73% contribution; see Figure 3B and File S2/Sheet4). The primary phenotypes contributing to Dimension 3 were also rather generic (e.g., increased compound excretion, decreased resistance to hyperosmotic stress, and absent or reduced utilization of nitrogen sources; see File S2/Sheet3), driven by *ade8* and *ade3* phenotypes (making a combined 73% contribution).

Overall, we conclude that the phenotypic classification of folate-dependent enzyme mutants is dominated primarily by the phenotypes of just two, *cdc21* and *gcv3*,with nearly all information regarding cell cycle progression arising from studies with *cdc21* mutants. Given the importance of 1C pathways in cell proliferation and the cell cycle-dependent changes in the levels of some of these enzymes we described above (Figure 2), we looked at their cell cycle-related phenotypes more closely.

**FIGURE 2.**
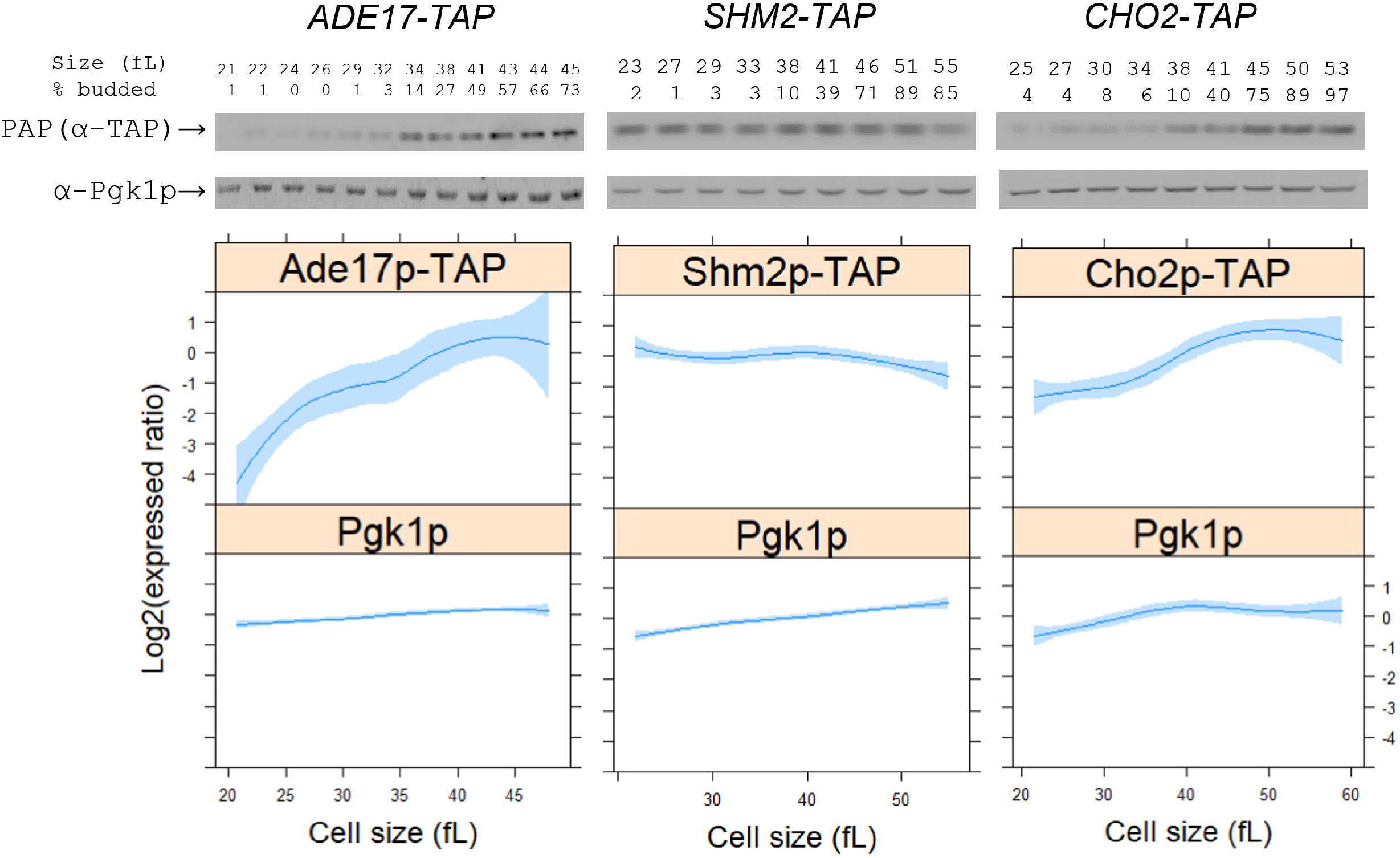
The levels of Ade17p, Shm2p, and Cho2p in the cell cycle. The abundance of TAP-tagged proteins was monitored in strains of the indicated genotype, as described in Materials and Methods. Samples were collected by elutriation in a rich, undefined medium (YPD) and allowed to progress synchronously in the cell cycle. Experiment□matched loading controls (measuring Pgk1p abundance) were also quantified and shown in parallel. (Top), representative immunoblots, along with the percentage of budded cells (% budded) and the cell size (in fL) for each sample. (Bottom), from at least three independent experiments in each case, the TAP and Pgk1p signal intensities were quantified as described in Materials and Methods. The Log2(expressed ratios) values are on the y-axis, and cell size values are on the x-axis. Loess curves and the standard errors at a 0.95 level are shown. All the immunoblots for this figure are in Figure S1, while the values used to generate the graphs are in File S1/Sheet1.

**FIGURE 3.**
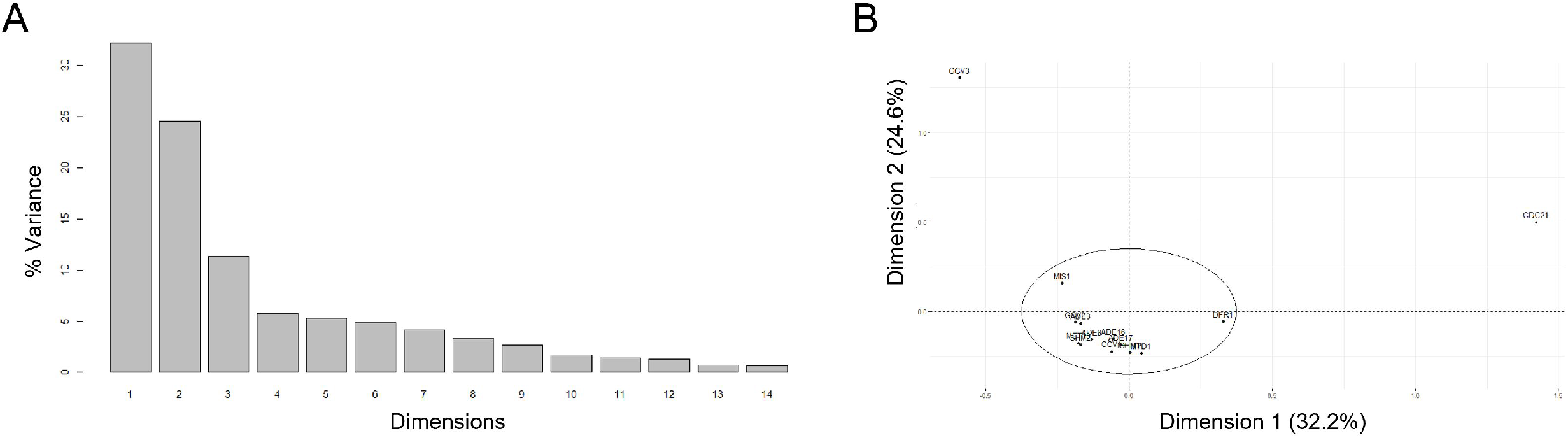
Multiple correspondence analysis of known phenotypes of mutants in folate-dependent enzymes. A, The sixty-six known phenotypes can be reduced to 14 dimensions, three of which explain 68% of the observed variance (y-axis). B, Plot of the individual gene contributions to the first two dimensions (x- and y-axes), which together explain >50% of the observed phenotypic variance, highlighting the role of *cdc21* and *gcv3* phenotypes in this classification. The contributions of the rest of the genes are grouped within the ellipse. The *FactoMiner* R language package was used in the analysis and generation of these plots. All the data are in File S2.

### Loss of Shm2p or Ade17p delays START

Since cell size changes are often accompanied by altered timing of cell cycle transitions, we first looked at the cell size distributions of homozygous diploid, asynchronous *ade17Δ* or *shm2Δ* cultures (Figure 4). Cells lacking Ade17p or Shm2p are born at normal size (Figure 4A, middle panel), but they have a larger mean size (Figure 4A, left panel), suggesting a delay at some later step in the cell cycle. Next, we measured the kinetics of cell cycle progression from synchronous, elutriated cultures. These experiments allowed us to determine the size at which half the cells in the culture are budded (a.k.a. critical size), marking passage through START in late G1, when cells commit to a new round of cell division and initiation of DNA replication. Loss of either Shm2p or Ade17p increased the critical size for START (Figure 4A, right panel). The cells grew larger before starting to bud compared to wildtype, consistent with the notion that they delay exiting from the G1 phase into the S. From the same experiments, we also measured the specific rate at which cells increase in size (k, Figure 4B), as a measure of cell growth. Cells lacking Shm2p have a lower k, indicating a growth defect. Hence, *shm2Δ* cells have a longer G1 phase because they have a larger critical size (Figure 4A, right) and reach that size at a slower rate (Figure 4B). On the other hand, *ade17Δ* cells increase in size at a normal rate (Figure 4B). These data point to a more specific role of Ade17p at START.

**FIGURE 4.**
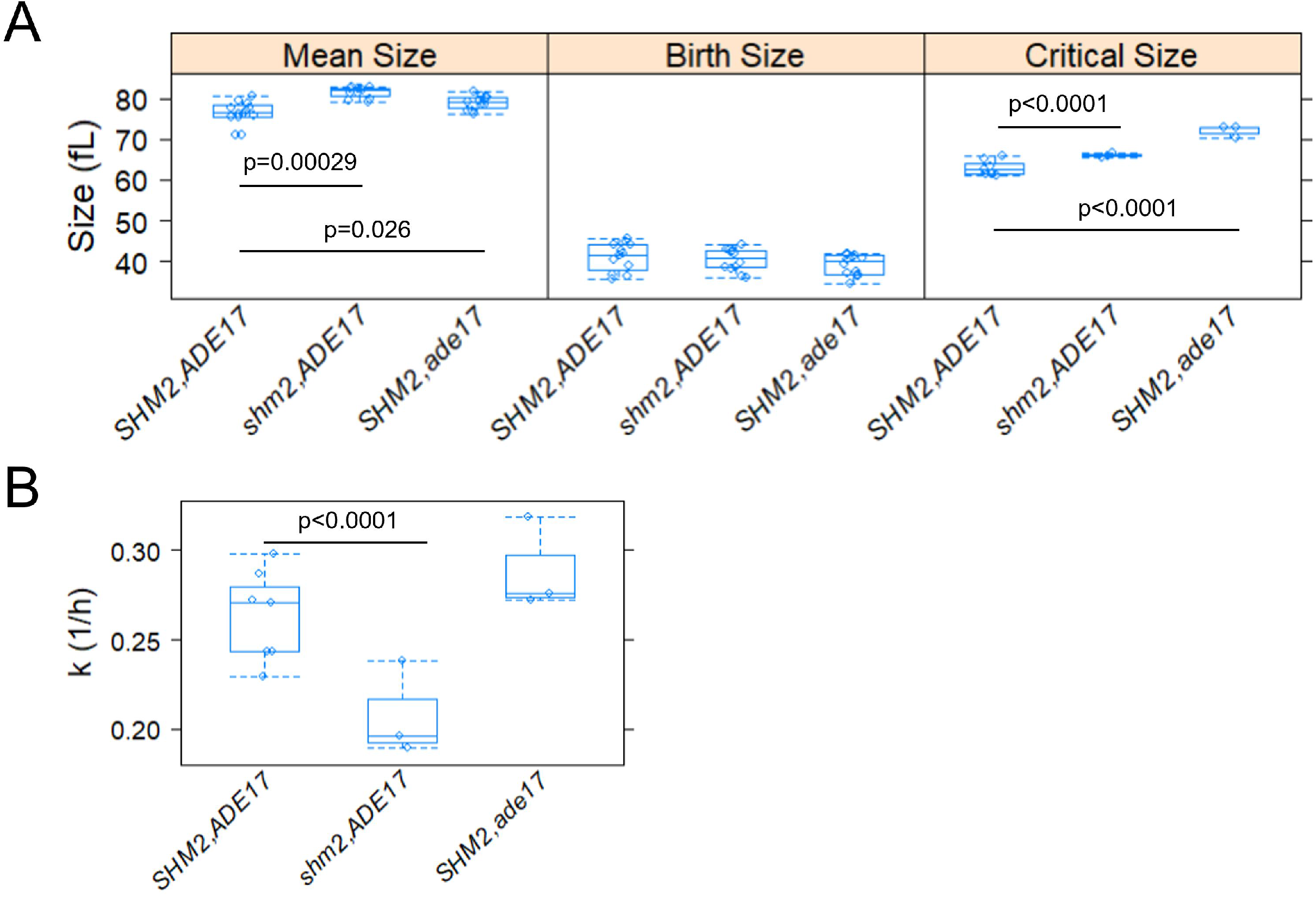
Loss of Shm2p or Ade17p delays START. A, Box plots showing the mean (left panel), birth (middle panel), and critical (right panel) size (y-axis) for the indicated strains. B, Box plots showing the specific rate of cell size increase (y-axis) for the indicated strains. All the values were calculated as described in Materials and Methods. Comparisons were made with the nonparametric Kruskal-Wallis rank sum test, and the indicated p-values calculated from the pairwise comparisons using the Wilcoxon rank sum test with continuity correction, using R language functions. The values used to generate the graphs are in File S1/Sheet2.

### *De novo* synthesis of phosphatidylcholine is required for cell growth and size control

Exogenous choline can be salvaged through the CDP-choline pathway to generate phosphatidylcholine (Dowd *et al*. 2001). Hence, to properly query the requirement for *CHO2* and *de novo* synthesis of phosphatidylcholine, we evaluated haploid *cho2Δ* cells not only in the standard, rich undefined medium (YPD) as above but also in synthetic minimal medium (SMM), without or with choline (at 1mM) supplementation. Asynchronous cells lacking Cho2p had a larger overall size in all media tested (Figure 5A, left panels), including in minimal media supplemented with choline (Figure 5A, left, middle panel), and they were also born slightly bigger (Figure 5A, middle panels).

**FIGURE 5.**
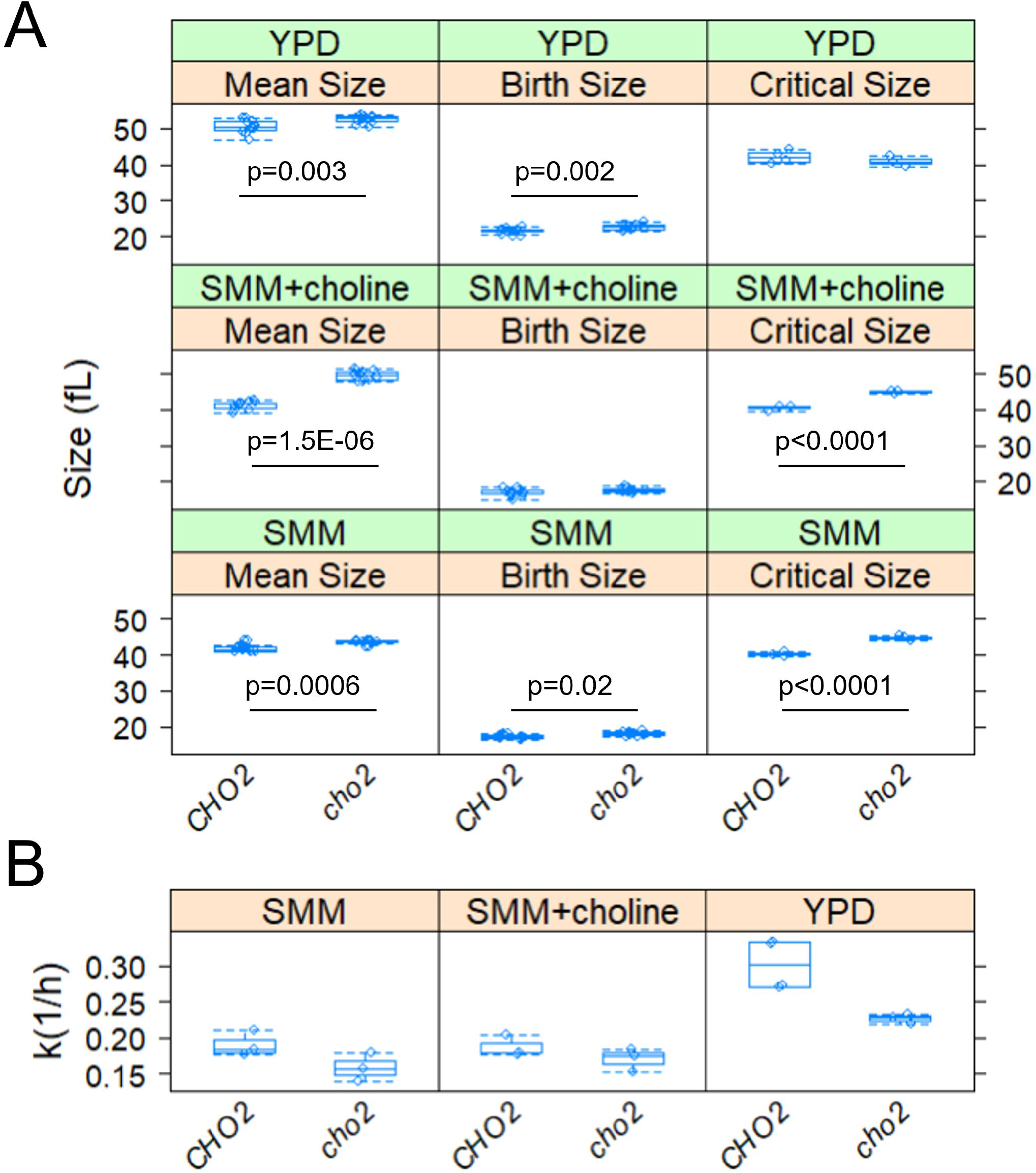
De novo synthesis of phosphatidylcholine is required for cell growth and size control at START. A, Box plots showing the mean (left panel), birth (middle panel), and critical (right panel) size (y-axis) for the indicated strains and medium. B, Box plots showing the specific rate of cell size increase (y-axis) for the indicated strains. All the parameters were calculated as described in the previous Figure. The values used to generate the graphs are in File S1/Sheet3.

To gauge cell cycle kinetics in all these conditions, we examined synchronous elutriated cultures, as described above (see Figure 4). Remarkably, *cho2Δ* cells had a significantly bigger critical size, albeit only in minimal media (Figure 5A, right panels). The latter effect was evident regardless of exogenous choline supplementation (Figure 5A, right, middle and bottom panels). Lastly, in all media tested, *cho2Δ* cells had a lower specific rate of size increase (Figure 5B), consistent with growth defects. These data suggest that Cho2p and the *de novo* pathway for PC synthesis are required for optimal cell growth and size homeostasis even when exogenous choline is available. To ensure that the budding index accurately reflects the position of *cho2Δ* cells in the cell cycle, in a separate elutriation experiment we also measured the DNA content of *CHO2^+^* and *cho2Δ* cells, in synthetic minimal media with or with exogenous supplementation with choline (Figure S2). The same conclusions were reached, that *cho2Δ* cells grow in size slower and they are larger when they initiate DNA replication.

### Loss of *de novo* PC synthesis dampens the kinetics of nuclear elongation late in the cell cycle

We had previously reported that increased lipogenesis promotes nuclear elongation and division (Maitra *et al*. 2022). To test if *de novo* synthesis of PC impinges on these late mitotic events, we analyzed the nuclear morphology of *cho2Δ* cells and their otherwise wild-type *CHO2* counterparts from synchronous elutriated cultures. Early G1 daughter cells were isolated as in Figure 5 and allowed to progress in the cell cycle in minimal media with or without choline. At regular intervals, the cells were fixed for immunofluorescence against the nucleoporin Nsp1p. Note that the nuclear envelope remains intact during mitosis in *S. cerevisiae*. Before metaphase, the nucleus elongates across the bud neck, between the mother cell and the large bud. The ratio of the long:short axes of the nucleus provides a metric of nuclear elongation (Maitra *et al*.2022). A ratio of one corresponds to a spherical nucleus, while values greater than 1 reflect nuclear elongation. In minimal media without choline, shortly after wild-type cells reach their critical size (~41fL, see Figure 6A, bottom), their nuclei start elongating, reaching their maximum length at ~46fL (Figure 6B, bottom). Cells lacking Cho2p have a larger critical size (~45fL, see Figure 6A, top), and their nuclei are maximally elongated at ~48fL (Figure 6B, top). However, instead of a well-formed, sharp peak, *cho2△* cells elongate their nuclei more gradually than wild-type cells (Figure 6B, top).

**FIGURE 6.**
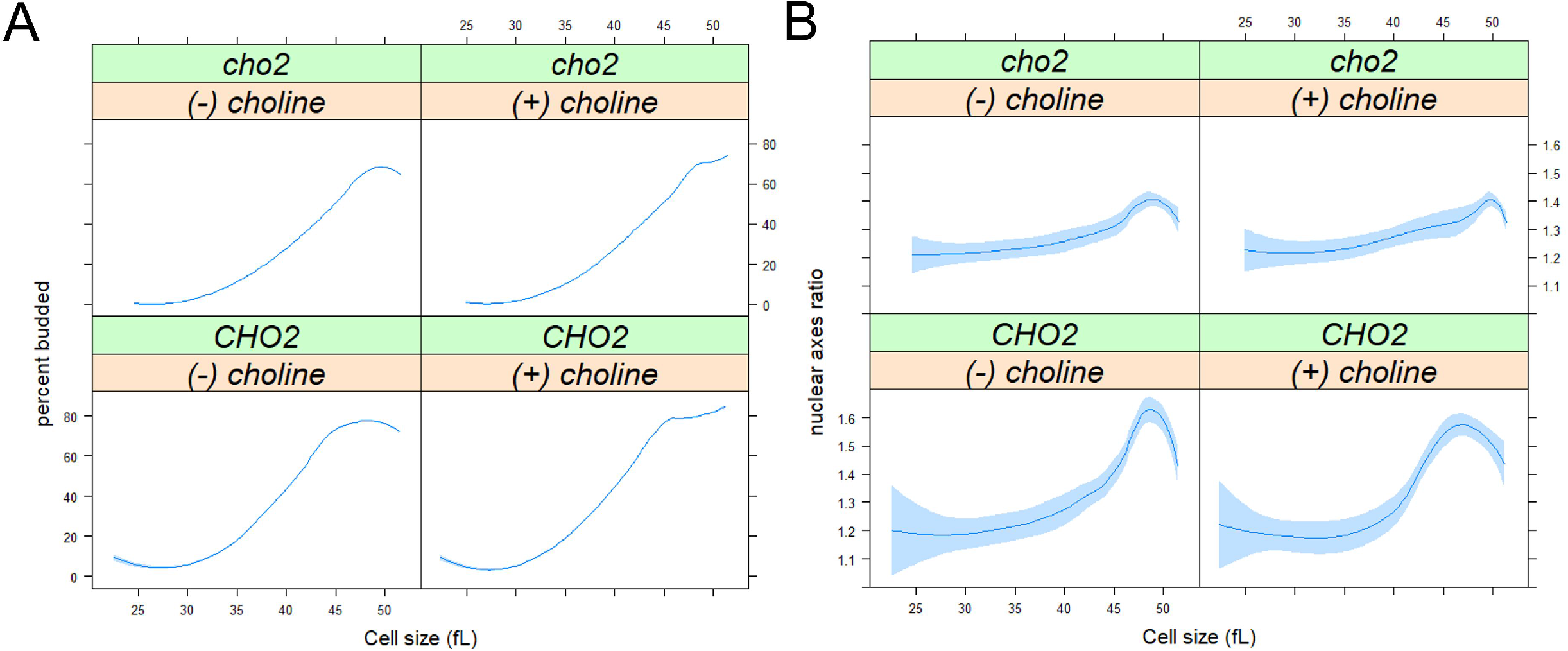
Loss of *de novo* phosphatidylcholine synthesis in cells lacking CHO2 dampens the kinetics of nuclear elongation late in the cell cycle. A, Synchronous cultures of the indicated strains and conditions were obtained by elutriation (see Materials and Methods), from which the percentage of budded cells (y-axis) is shown against the mean cell size (in fL; x-axis). B, cells were processed for fluorescence microscopy to visualize the nucleus from the same samples as in A, as described in Materials and Methods. Nuclear axes ratio (y-axis) is shown against the mean cell size (in fL; x-axis), from >2,000 cells/strain. Loess curves and the std errors at a 0.95 level are shown. The values used to generate the graphs are in File S1/Sheet4.

Choline supplementation did not suppress this behavior (if anything, it might have delayed somewhat the full elongation of the nuclei of *cho2Δ* cells; compare the top panels in Figure 6B). These results suggest that *cho2Δ* cells elongate their nuclei less coherently or synchronously than wild-type cells, implicating the *de novo* pathway of PC synthesis in nuclear elongation and division.

### Metabolic profiling of cells lacking Ade17p, Shm2p, or Cho2p

Since 1C enzymes control several key metabolic outputs (Figure 1), we sought to place the cell cycle phenotypes we uncovered in the context of possible metabolic changes. First, we asked if there was any change in histone methylation. One might expect that reduced PC synthesis in *cho2Δ* cells might lead to increased use of 1C units for histone methylation (Ye *et al*. 2017). In the same strains and media we described above (see Figures 4,5), we measured the levels of trimethylated histone H3 on Lys4 (H3K4me3) compared to the total levels of histone H3 (see Materials and Methods). We found no significant changes (p>0.05 in the Kruskal-Wallis test) in the ratio of H3K4me3:H3 levels among the different media or strains (*ade17Δ, shm2Δ*, or *cho2Δ*cells) we examined (Figure S3).

To gain a broader overview of metabolic alterations in the mutants we examined, we measured the distribution of the steady-state levels of metabolites and lipids by mass spectrometry (see Materials and Methods). Different analytical platforms were used to separate and identify primary metabolites (File S3), biogenic amines (File S4), and complex lipids (File S5). We combined the identified primary metabolite and biogenic amine datasets, while the complex lipid datasets were kept separate. To identify metabolite and metabolic pathway enrichment, we used the MetaboAnalyst platform (Chong *et al*. 2019). Pairwise comparisons of the metabolome between wild type vs. *ade17Δ* and wild type vs. *shm2Δ* cells growing in undefined, rich YPD medium revealed few significant changes (Figure 7A, top), and these changes were very similar in cells lacking Ade17p or Shm2p. No group was significantly enriched among the metabolites with lower levels (1.5-fold change and p<0.05) in *ade17Δ* or *shm2Δ* cells. On the other hand, among metabolites with significantly higher levels in *shm2Δ* cells (19 compounds), the only group significantly enriched were metabolites associated with gluconeogenesis (p=0.0458 with Holm–Bonferroni correction). The same trend was evident in *ade17Δ* cells (p=0.0646 with Holm–Bonferroni correction from the 23 metabolites with significantly higher levels). Although there were few metabolite changes associated with loss of either Shm2p or Ade17p (Figure 7A, top), these mutants seem to acquire gluconeogenic signatures, even though they were growing in a glucose-replete, rich medium. Lastly, while in both *shm2Δ* and *ade17Δ* cells there was a reduction in the levels of some complex lipids compared to wild-type cells (Figure S6A, top), the lipids that changed in abundance were different. Compared to wild-type, *shm2Δ* cells had lower levels of triacylglycerols (p=1.79E-05, with Holm-Bonferroni correction). On the other hand, *ade17Δ* cells had lower levels of lyso-PC (p=4.22E-10, with Holm-Bonferroni correction), PC (p=3.98E-06, with Holm-Bonferroni correction), and monoacylglycerophosphocholines (p=0.0124, with Holm-Bonferroni correction). These data suggest that the loss of Shm2p reduces triacylglycerols, while the loss of Ade17p may direct lipid synthesis away from PC, resembling qualitatively cells lacking Cho2p, as described below. Nonetheless, despite these statistically significant differences, we note that the overall number of lipid species with altered levels and their fold-change in abundance in *ade17Δ* and *shm2Δ* cells compared to wild-type cells was relatively small (Figure S6A, top).

**FIGURE 7.**
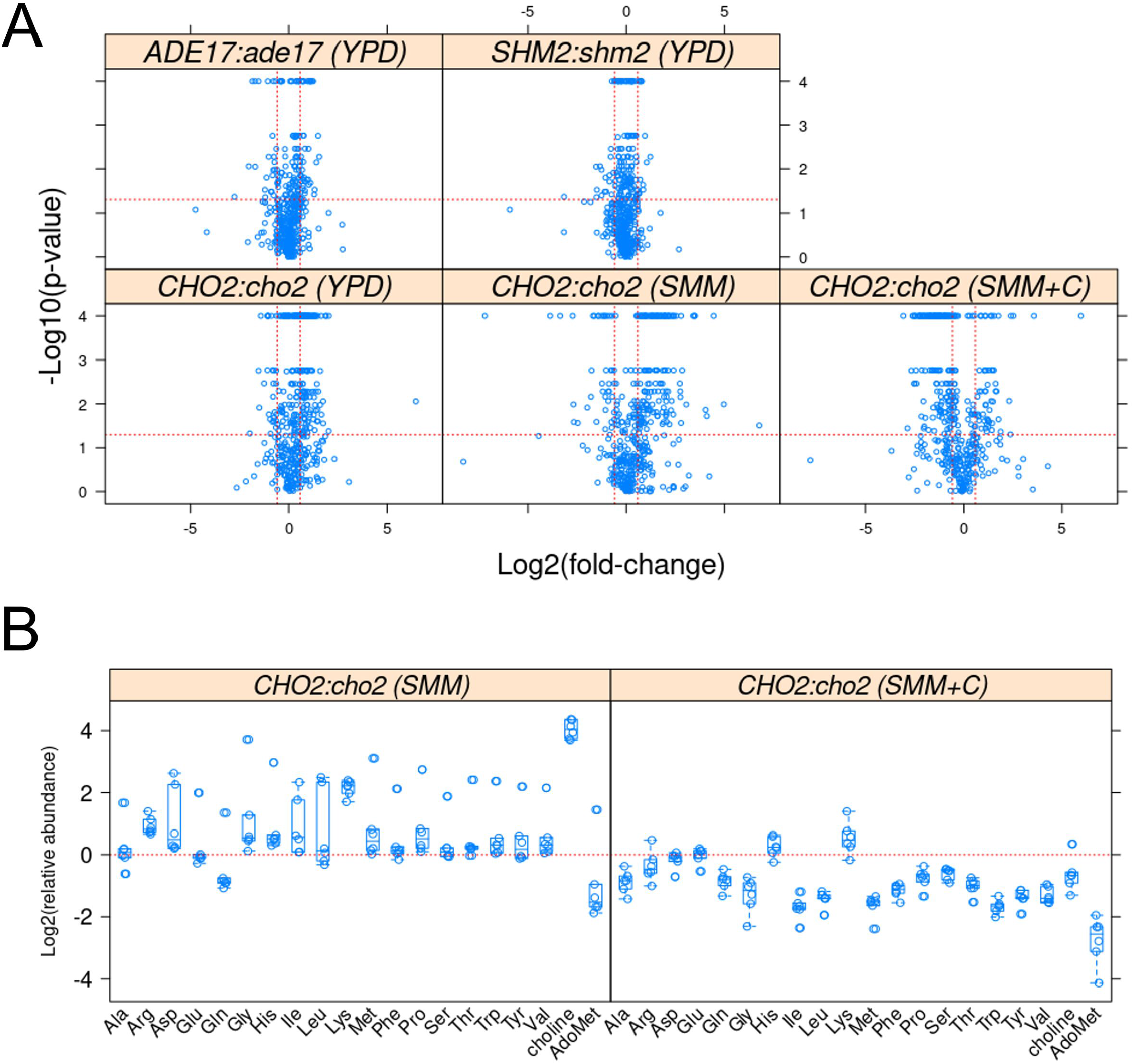
Changes in primary and biogenic amine metabolites in cells lacking Adel7p, Shm2p, or Cho2p. A, Metabolites whose levels changed in the indicated pairwise comparisons and media (shown in parenthesis above each panel) were identified from the magnitude of the difference (x-axis; Log2-fold change) and statistical significance (y-axis), indicated by the red lines. The analytical and statistical approaches are described in Materials and Methods. The values used to generate the graphs are in File S1/Sheet6. Note that the lowest calculated p-values from the robust ANOVA were at the 0.0001 level. B, Boxplots of the levels of the amino acids (x-axis) detected in the biogenic amine dataset (see Materials and Methods), shown as Log2-transformed relative abundance between wild type (*CHO2*) and *cho2Δ* cells (y-axis), from six independent samples in each case. The levels of choline and AdoMet (S-adenosylmethionine) from the same measurements are also shown. The values used to generate the graphs are in File S1/Sheet7.

The metabolite changes were profound in *CHO2* vs. *cho2Δ* cells, with >200 primary metabolites and biogenic amines displaying significantly altered steady-state levels (see Figure 7A, bottom), in all three different media we tested (YPD, SMM, SMM+choline). The compounds with reduced levels in *cho2Δ* cells were significantly enriched for groups associated with amino acid metabolism and the overall pathway of “aminoacyl-tRNA biosynthesis” (p=0.0049 with Holm-Bonferroni correction). In other words, amino acid pools were depleted in *cho2Δ* cells. The same pattern was observed in the minimal (SMM) medium, comparing *CHO2* vs. *cho2Δ* (p=0.0321 with Holm-Bonferroni correction; see Figure 7B, left panel). Strikingly, comparing the metabolite repertoire in wild-type cells in YPD vs. SMM media (see Figure S5A), which queries nutrient effects, metabolites associated with the overall pathway of “aminoacyl-tRNA biosynthesis” were also significantly enriched (p=8.08E-05 with Holm–Bonferroni correction). Intracellular amino acid levels were higher in wild-type cells cultured in the rich YPD medium vs. the minimal medium (Figure S5B, left panel). These results suggest that loss of *de novo* PC synthesis in *cho2Δ* cells in rich undefined media leads to metabolic changes typically associated with nutrient shifts to poorer, minimal media. We note that, as expected, choline levels were dramatically reduced (~25-fold) in *cho2Δ*cells in the minimal SMM media (Figure 7B, left panel). But the methylation potential of these cells remained high, having elevated AdoMet (S-adenosylmethionine) levels (Figure 7B, left panel), consistent with the unchanged histone methylation levels we reported above (Figure S3). Regarding changes in the lipidome, in both YPD and SMM media, loss of Cho2p significantly reduces the levels of >60 complex lipids (Figure S6A, bottom panels). As expected, *cho2Δ* cells had lower levels of lyso-PC, PC, and diacylglycerophosphocholines (p<3E-06, with Holm–Bonferroni correction).

These conclusions in *CHO2* vs. *cho2Δ* cells were reinforced with the metabolite measurements in the minimal SMM medium supplemented with 1mM choline. Wild-type cells showed no significant changes (see Figure S5A, left panel). But the changes in *cho2Δ* cells in SMM medium were reversed by adding 1mM choline. The pathway of “aminoacyl-tRNA biosynthesis” became significantly enriched in *cho2Δ* cells compared to wild type (p=3.18E-06 with Holm–Bonferroni correction), as reflected in their higher amino acid levels (Figure 7B, left panel). In wild-type cells, addition of choline did not cause any changes in amino acid levels (Figure 5SB, right panel). Taken together, these results strongly suggest that inability to carry out *de novo* PC synthesis in *cho2Δ*cells leads to profound changes in amino acid homeostasis. Lasty, the lipid changes mentioned above between wild-type and *cho2Δ* cells were also completely reversed by supplementation with 1mM choline (Figure S6A, bottom right panel), with cells lacking Cho2p even having higher levels of lyso-PC and diacylglycerophosphocholines (p<0.012, with Holm–Bonferroni correction).

## DISCUSSION

Our data reveal that the levels of some 1C enzymes are dynamic in the cell cycle, and the corresponding loss-of function mutations lead to distinct cell cycle phenotypes and metabolic changes. We place these findings in the context of the existing literature and discuss possible implications.

In humans, loss-of-function mutations have been described for ATIC (*ADE17*ortholog), SHMT1 (*SHM2* ortholog), and PEMT (*CHO2* ortholog). Patients with ATIC mutations accumulate AICAR (5-aminoimidazole-4-carboxamide-1-β-D-ribofuranoside), and display a wide range of developmental abnormalities (Ramond *et al*. 2020). At the cellular level, in patient-derived fibroblasts, a reduced ability to form purinosomes and drive purine biosynthesis correlates with clinical phenotypes of individual patients (Baresova *et al*. 2012). Interestingly, however, moderately reducing ATIC activity in adult mice with a specific inhibitor is beneficial, leading to an increase in AICAR and AMP-activated kinase (AMPK) activity, ameliorating metabolic syndrome phenotypes (Asby *et al*. 2015). How does one reconcile these observations? It is reasonable to expect that during embryogenesis and early adult life, when cell proliferation underpins developmental milestones in animals, loss-of-function perturbations in 1C metabolism would lead to profound phenotypes, but perhaps not so later in life. This is also the basis for the selective effects of antifolates in chemotherapy. Indeed, the LD_50_ for methotrexate given to 5-week-old mice is 59mg/kg, whereas that for 16-week-old mice is 284mg/kg (Freeman-Narrod and Narrod 1977). In yeast, perhaps analogous to a proliferating animal tissue, complete loss of ATIC, in double *ade16,17Δ* mutants, leads to inviability unless the cells are supplemented with exogenous adenine (Tibbetts and Appling 2000). The single *ade17Δ* cells we evaluated here allowed us to reveal rather specific effects at START (Figure 4). Even though the growth rate is unaffected, the larger critical size of *ade17Δ* mutants is consistent with the interpretation that cells may monitor some aspect of purine biosynthesis before committing to a new round of cell division. In this regard, having the levels of Ade17p under translational control, rising as cells exit G1 (Figure 2), provides another way for the cell to link protein synthesis and overall biosynthetic capacity with metabolic pathways needed for cell division, contributing to the general task of coupling cell growth with cell division.

There are no reported SHMT1 mutations causing human pathologies. In mice, however, *Shmt1* knockout animals are a genetic model of folate deficiencies (MacFarlane *et al*. 2008). SHMT1 has been shown to directly interfere with folate metabolism and its loss leads to folate deficiency, sensitizing embryos to neural tube defects (Beaudin *et al*. 2011). Adult *Shmt1^-/-^* mice, however, are fertile and healthy. Their methionine plasma levels were lower (at 43μM), irrespective of whether they were on a folate-replete or folate-limited diet (MacFarlane *et al*. 2008). Wild-type mice fed a folate/choline-deficient diet for five weeks had reduced methionine plasma levels too (by ~25%, from 60μM to 45μM), while serine and glycine levels did not change (MacFarlane *et al*. 2008). Yet, significant Shmt1-dependent changes to methylation capacity, gene expression, and purine synthesis were not observed (Macfarlane *et al*. 2011). Overall, it appears that loss of the cytoplasmic serine hydroxymethyltransferase in mammals leads to minimal, if any, adverse phenotypes in adults. In the constantly dividing *S. cerevisiae*cells, on the other hand, the lower growth rate and larger critical size of *shm2Δ* cells (Figure 4) likely reflect the dependency on cytoplasmic hydroxymethyltransferase for optimal cell proliferation.

Mammals rely mainly on the salvage PC synthesis pathway, incorporating dietary choline into PC, except for the liver, where Pemt catalyzes significant *de novo* PC synthesis (Walkey *et al*. 1997). The corresponding knockout mice were normal, including in their hepatocyte morphology, bile composition, or plasma lipid levels (Walkey *et al*. 1997), arguing for minimal contributions, if any, for the *de novo* PC pathway in mammals. In yeast, Cki1p encodes choline kinase, which catalyzes the first step in phosphatidylcholine synthesis in the choline salvage pathway. We previously showed that *cki1Δ* cells have a slightly smaller critical size in rich, YPD medium (Maitra *et al*. 2022). Here, we found that *cho2Δ* cells have a significantly larger critical size in minimal media, regardless of whether exogenous choline becomes available (Figure 5). They also have a reduced specific rate of size increase in all media, even in YPD (Figure 5B), suggestive of rather broad impacts on cellular physiology. Overall, unlike the situation in animal cells, our results point to important roles in proliferating *S. cerevisiae* cells for *de novo* PC synthesis. Our data suggest that nuclear elongation late in the cell cycle is one such role.

Nuclear division places heavy demands for new membrane material in the relatively short time it takes for mitosis. This is true not only in fungi, which undergo closed mitosis but in animal cells as well, when the fragmented nuclear membrane must reform around each of the nuclei in telophase. In *S. pombe* and *S. cerevisiae*, loss-of-function mutations in lipid synthesis change the size and shape of the nucleus (Santos-Rosa *et al*. 2005; Witkin *et al*. 2012; Walters *et al*. 2012; Siniossoglou 2013; Kume *et al*. 2017; Zach and Prevorovsky 2018). In animal and yeast cells, inactivating acetyl-CoA carboxylase, the rate-limiting lipogenic enzyme, blocks cells in mitosis, and exogenous fatty acids cannot rescue the cell cycle arrest (Schneiter *et al*. 1996; Al-Feel *et al*. 2003; Scaglia *et al*. 2014). Conversely, we have shown that increased lipogenesis promotes nuclear division (Maitra *et al*. 2022). In this context, it might appear unsurprising that disrupting PC synthesis in *cho2Δ* cells alters the kinetics of nuclear elongation (Figure 6). However, we also showed that nuclear elongation kinetics were unaffected in *cki1Δ*cells with impaired PC synthesis from the salvage pathway (Maitra *et al*. 2022). In contrast, the data in this report highlight the need for *de novo* PC synthesis during nuclear division. Our data also revealed unexpected metabolome changes in cells lacking Cho2p (Figure 7). To our knowledge, this is the first time that such a link between *de novo* PC synthesis and amino acid levels has been reported. Astonishingly, even in rich media, *cho2Δ* cells behave as if they are in minimal media (Figures 7 and S5). How the inability to synthesize PC *de novo* also leads to a lowering of amino acid levels is unclear.

In summary, the presented results revealed unappreciated contributions of 1C metabolism to cell cycle progression. 1C metabolism is an excellent platform for integrating various metabolic inputs and outputs with cell division. This study adds to effort to decipher how specific cell cycle processes are linked to the enzymes of 1C metabolism. The results we presented could also inform research that may not be focused on cell cycle progression, especially in the context of several recent studies linking 1C metabolism with longevity mechanisms in multiple systems (Maitra *et al*.2020; Enriquez-Hesles *et al*. 2021; Annibal *et al*. 2021; Lionaki *et al*. 2022).

## DATA AVAILABILITY

Strains and plasmids are available upon request. The authors affirm that all data necessary for confirming the conclusions of the article are present within the article, figures, and tables. The Supplementary Material is available through figshare (https://doi.org/10.6084/m9.figshare.21390978.v1), and includes the following:

Reagent Table
Figure S1: Source immunoblots for the protein abundances of Figure 2.
Figure S2: DNA content profiles of synchronous, elutriated *cho2Δ*cells.
Figure S3: Quantification of histone methylation levels.
Figure S4: Source immunoblots for histone methylation levels.
Figure S5: Comparison of metabolite levels in haploid wild type cells cultured in different media.
Figure S6: Comparison of complex lipid levels in different strains and media.
File S1:Source data for the graphs in Figures 2, 4, 5, 6, 7, S3, S5.
File S2:File with all the loss-of-function phenotypic analyses reported in the literature for genes encoding folate-dependent enzymes and the multiple correspondence analysis (MCA) we performed.
File S3:File with the raw mass spectrometry data for primary metabolites.
File S4:File with the raw mass spectrometry data for biogenic amines.
File S5:File with the raw mass spectrometry data for complex lipids.

## ACKNOWLEDGEMENTS

We thank the UC Davis West Coast Metabolomics Center for their help with metabolite measurements.

